# Integrating multiple sensory modalities during dyadic interactions drives self-other differentiation at the behavioral and electrocortical level

**DOI:** 10.1101/2025.08.01.668120

**Authors:** Ugo Giulio Pesci, Giovanna Cuomo, Vanessa Era, Matteo Candidi

**Affiliations:** Department of Psychology, Sapienza University, Rome, Italy; IRCCS Fondazione Santa Lucia, Rome, Italy

**Keywords:** interpersonal interactions, sensorimotor coordination, sensory processing, cross-modal processing

## Abstract

Interpersonal motor interactions represent a key setting for processing signals from multiple sensory channels simultaneously, possibly modulating cross-modal multisensory integration, that is a crucial perceptual mechanism where different sensory sources of information are combined into one single percept. Here we explored whether integrating sensorimotor signals while interacting with a partner can lead to shared sensorimotor representations, and possibly to a recalibration of individual’s multisensory perception. In detail, we investigated whether engaging individuals in dyadic activities that utilized either single (e.g., visual or tactile/proprioceptive) or combined (e.g., visuo-tactile/proprioceptive) sensory modalities would impact the behavioral and electrocortical markers associated with interpersonal cross-modal integration. We show that interactions requiring the integration of multiple sensory modalities lead to higher interpersonal differentiation resulting in reduced interpersonal cross-modal integration in a subsequent spatial detection task and alter its distributed neural representations. Specifically, the neural patterns elicited by interpersonal visuo-tactile stimuli vary based on the sensory nature of the previous interpersonal interaction, with the one involving multiple sensory modalities resulting in improved performance of a neural classifier. These findings suggest new avenues for sensorimotor approaches in social neuroscience, emphasizing the malleability of self-other representations based on the nature of interpersonal interactions.

**SIGNIFICANCE STATEMENT:** Successful social interaction relies on the dynamic integration of sensory and motor signals between individuals. Here, we show that the sensory channels engaged during interpersonal coordination recalibrate how the brain processes multisensory information about the self and others after having interacted with them. Specifically, interactions involving integrated visual and proprioceptive feedback sharpen self-other distinction and reduce cross-modal interference. These effects manifest in both behavioral responses and early neural activity patterns, revealing plastic, offline changes in multisensory integration shaped by social sensorimotor contingencies. Our findings highlight the role of interpersonal experience in shaping the functional responses to multisensory stimuli and provide novel insights into the sensorimotor foundations of social embodied cognition.

## INTRODUCTION

Sensorimotor integration (SMI) refers to the process by which sensory events are used to program, and are combined with, one’s motor commands. Modern conceptions (Wolpert et al., 1995; Körding & Wolpert, 2004) hold that not only perception guides action, but that through predictive mechanisms, sensory perception is continuously refined by one’s movements (Egger et al., 2019; for a review on predictive mechanisms during movement control see Latash, 2021; in vision for action see Fiehler et al., 2019; in the tactile domain see Fuehrer et al., 2022). A key aspect of SMI involves the acquisition of sensorimotor contingencies (SMCs)— learned associations between movements and the resulting sensory changes (O’Regan & Noë, 2001). Originally considered within individual perception and action, SMCs have more recently been proposed as central to interpersonal coordination, and the framework of social sensorimotor contingencies (socSMCs) has been proposed to study how individuals integrate sensory signals from others with their own behavior (Lübbert et al., 2021). Social SMCs are thought as dynamic informational and sensorimotor coupling across agents which can mediate the deployment of action-effect contingencies in social contexts. Such mechanisms may underpin interpersonal learning during development (Goupil et al., 2024) and support more complex social functions like mentalizing (Tamir & Thornton, 2018; Kingsbury & Hong, 2020).

SMI, particularly in the context of motor control, can be seen as a special case of multisensory integration (MSI) (Aschersleben & Bertelson, 2003; Repp, 2005; Stekelenburg et al., 2011; Jagini, 2021), and this is particularly relevant during interpersonal interactions, where sensory feedback depends on both one’s own and another’s movements (Pezzulo et al., 2013; Jagini, 2021). These scenarios demand the concurrent and dynamic processing of signals from multiple modalities, possibly giving rise to interpersonal cross-modal effects. MSI involves combining inputs from different sensory channels in a statistically optimal way (Ernst et al., 2002; Fetsch et al, 2011; Knill & Saunders, 2003), allowing for more reliable perception and for solving the so-called “causal inference problem” – i.e., determining whether inputs from different modalities arise from the same source (Körding et al., 2007; Rohe et al., 2019). MSI also contributes to maintaining a stable yet flexible representation of the body, crucial not only for interacting with the environment (Blanke et al., 2015), but also for distinguishing between events affecting oneself versus another person (e.g., Fossataro et al., 2022).

Importantly, sensory processing is modulated during movement via mechanisms like gating and attenuation (Rushton et al., 1981; Starr & Cohen, 1985; Seki & Fetz, 2012; Kilteni & Ehrsson, 2022; Fuehrer et al., 2022; Arikan et al., 2023). Yet, how MSI operates during or after interpersonal motor interactions—particularly when multiple sensory modalities are involved—remains poorly understood. These interactions may lead to plastic changes in how both self- and other-related information is perceived and represented—what we refer to as interpersonal integration effects.

To investigate the sensorimotor basis of these effects, we examined whether engaging in interpersonal activities involving either single (e.g., visual or tactile/proprioceptive) or combined (e.g., visuo-tactile/proprioceptive) sensory modalities alters the behavioral and neural markers of interpersonal cross-modal integration. Specifically, we expected to (1) replicate and extend previous findings of cross-modal integration near both one’s own and another person’s body (Maravita et al., 2002; Fossataro et al., 2022); and (2) observe that multimodal interpersonal interaction leads to a recalibration of interpersonal representations possibly based on multisensory behavior principles, such as processes related to the perception of cross-modal (in)congruencies between stimuli (Spence et al., 1998; Pavani et al., 2000; Spence et al., 2004; Otto et al., 2013) delivered on/around the interactors’ hands. In detail, we hypothesized either 1) that interacting with another individual leads to an “integration” of her body schema into one’s own, thus leading to a stronger interpersonal cross-modal interference (i.e., stronger incongruency effect), or 2) that interactions requiring functional integration across modalities would enhance the reliability of ones’ own, the other’s, or both agents’ body schema, leading to a better distinction of the other, thereby reducing interpersonal cross-modal interference (i.e., weaker incongruency effects). Regarding the neural responses elicited by visuo-tactile stimuli, we expected to find a modulation of the patterns related to cross-modal incongruency processing, as well as new time-resolved signatures of a recalibration of visuo-tactile perception following different interpersonal tasks.

## MATERIALS AND METHODS

### Participants

Twenty-five subjects (15 males) with any neurological or psychiatric condition reported were involved in the study. The sample size was determined through a prospective power analysis performed with the software More Power (Campbell and Thompson, 2012). We inserted as expected effect size the partial eta squared value (0.22) observed in Fossataro et al., 2022, where a similar cross-modal interpersonal set-up was deployed. The analysis indicated that a 3 × 2 × 2 within-subject design, a power of 0.90 and a partial eta squared value of 0.22, required a sample size of at least 24 participants. Participants’ age varied between 20-29 years (mean age = 24.8 ± 2). All participants were right-handed with normal or corrected-to-normal vision. Participants were naive as to the aim of the experiment and were informed of the purpose of the study only after all the experimental procedures were completed. The experimental protocol was approved by the ethics committee of the Fondazione Santa Lucia (Rome, Italy) and was carried out in accordance with the ethical standards of the 1964 Declaration of Helsinki and later amendments.

### Stimuli of the visuo-tactile task

The experimental set-up that indexes intersensory effects incorporates and adapts methodologies adopted by previous studies investigating visuo-tactile integration processes by means of EEG (Sciortino et al., 2022; Fossataro et al., 2022). Specifically, participants sat in front of a desk where a 1024 x 768 resolution LCD monitor laid horizontally at ∼60 cm from their eyes. During the MSI task (see below), subjects were asked to place their left-hand palm down on the screen (11 cm to the left of their body midline), right (∼15 cm) to the confederate’s left hand, and to constantly fixate a white cross (1 cm diameter) positioned between the confederate’s index and thumb while their own hand was covered by a dark blanket and hidden from view. (Fig. 1A). Tactile stimuli were delivered to the participant’s left index/thumb using two 12 V round solenoids (0.8 cm in diameter) driving a magnet (1.5 mm in diameter) to the finger pad. Every time a current was delivered to the solenoid the magnet contacted the finger. The delivery of the current, and thus the presentation of the tactile stimuli, was controlled by a custom-built tactile controller (Heijo Research Electronics), which in turn was connected to the stimulus presentation computer. Although mechanical stimuli are less controllable than electrical ones in term of duration, each tactile stimulus was ∼5 ms long. Before the beginning of the experimental procedure, subjects were explicitly asked if they could clearly feel the tactile stimuli. Visual stimuli consisted of flashes (1.2 cm in diameter) occurring next to the tip of the confederate’s index/thumb. Visual and tactile stimuli were always presented simultaneously. Paradigm presentation and subjects’ response were controlled and recorded using E-Prime 2.0 Professional software (Psychology Software Tools Inc., Pittsburgh, PA).

**Figure 1.**
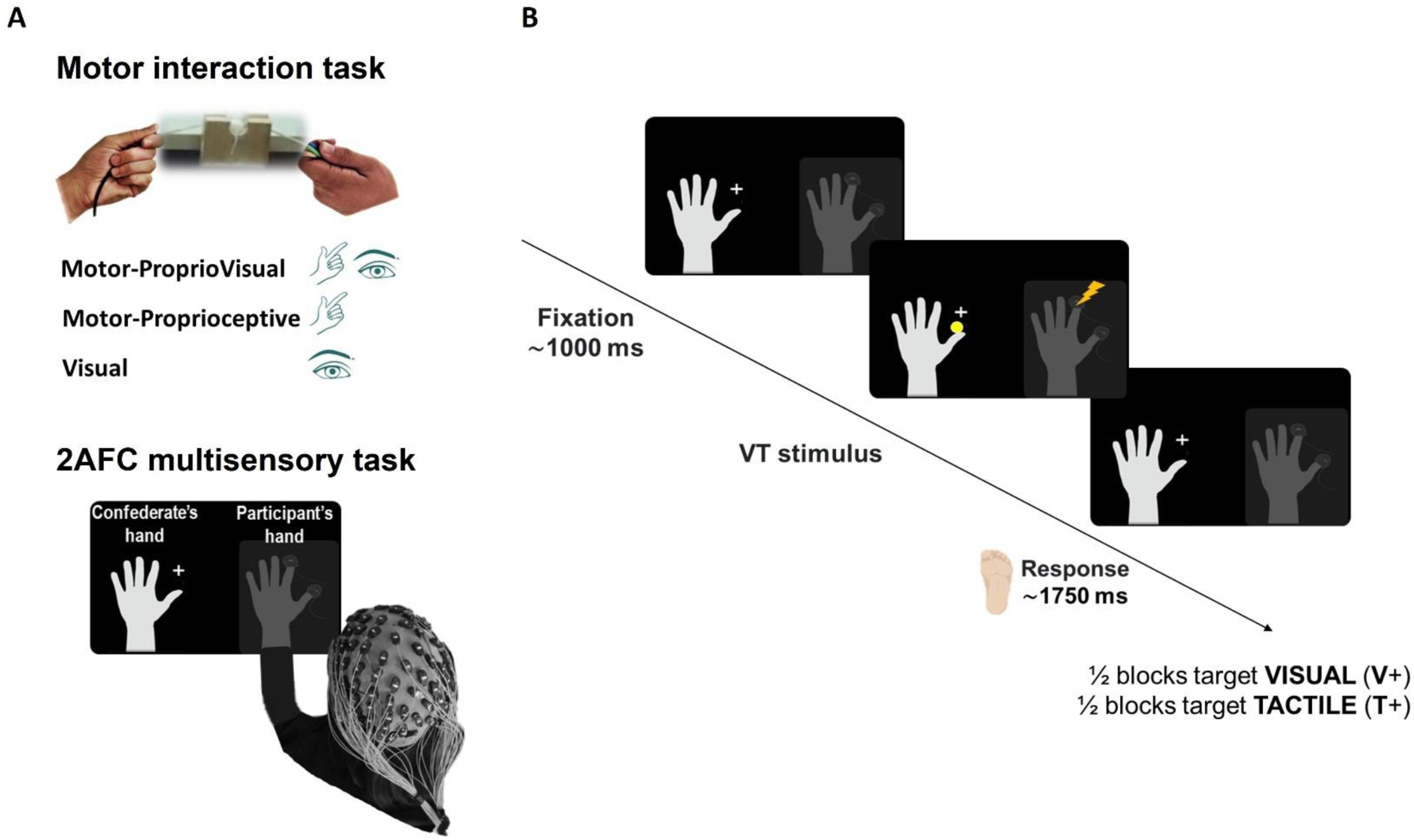
A) Experimental set-up. Graphical representation of the set-up during the interpersonal motor interaction task (upper plot), and the 2AFC interpersonal visuo-tactile task (bottom plot). B) Example timeline of a trial in the 2AFC multisensory task. Participants received tactile stimuli on their left hand (covered from view), either on the thumb or on the index, while visual flashes synchronously appeared close to the thumb or the index of the partner’s left hand laying on a screen next to them. Subjects had to report the location of either the tactile (T+) or of the visual (V+) stimulus as fast and as accurately as possible, in no more than 1750 ms. Each trial began with the fixation cross on the screen for 1000 ms (jittered from 750 to 1250 ms).

### Experimental design of the interpersonal interaction task

Participants’ visuo-tactile intersensory effects were measured after they had experienced three different behavioral tasks that we label Interpersonal Interaction Task. In detail, individuals were asked to observe or interact with a partner’s movements by 1) passively observing them (Visual condition - V), 2) moving together with them with their eyes closed (Motor-Proprioceptive condition – M-P), or 3) moving together with them with their eyes open (Motor-ProprioVisual condition – M-PV). Specifically, they performed a candle-sawing activity (Soliman et al., 2015), that required the participant and the confederate each to use their left hand to bimanually and rhythmically operate a cutting wire sideways to saw a candle positioned horizontally on a custom-made holder. Importantly, being the wire flexible, such task required the participants to coordinate their movement in space (pulling the wire) and time (following and pacing the couple’s movement rhythm) to reach the goal of sawing the candle. Moreover, such set-up allowed to isolate the motor-tactile/proprioceptive channel (when subjects had to perform the task with their eyes closed – Motor-Proprioceptive condition), the visual channel (when subjects had to passively observe the movement of the confederate – Visual condition), as well as to combine the two (when subjects had to synchronize their movement while observing the confederate’s hand – Motor-ProprioVisual condition). After the interaction, cross-modal interpersonal integration processes were measured by means of visuo-tactile interference effects realized on one’s own and the partner’s hand (similar to Maravita et al., 2003 and Heed et al., 2010). Specifically, participants received tactile stimuli on their left hand (the same one they used during the interaction), either on the thumb or on the index, while the visual stimuli synchronously appeared close to the thumb or the index of the partner’s left hand laying on a screen next to them (Fig 1B). Importantly, during this task the participant’s hand was covered with a dark blanket and hidden from view. White noise was played through headphones during the entire task to attenuate EEG artifacts due to the noise produced by the mechanical tactile stimulators. In half of the trials, the location of the visual and tactile stimuli differed (i.e. a flash appeared close to the confederate’s index while the participants received a tactile stimulus on his/her thumb – Congruency factor). Subjects were instructed to press a pedal either with their right or left foot, depending on the location of the target stimulus (left for stimuli on/close to the index and right for stimuli on/close to the thumb), in no more than 1750 ms. Moreover, to test the role of top-down effects on multisensory integration processes, subjects would be asked to give their response based on the location of either the tactile stimulus (Tactile Relevance – T+) in half of the trials, or the visual stimulus (Visual Relevance – V+ - Task Relevance factor) in the other half of the trials. The inter-trial-interval (ITI) was of 1000 ms (jittered from 750 to 1250 ms). For each interaction (M-PV, M-P, V), 4 consecutive experimental blocks were run. Each block began with an interactive session lasting 3 minutes, followed by a multisensory integration (MSI) measurement consisting of 40 trials, for a total of 160 trials per interaction (40 T+-Congruent trials, 40 T+-Incongruent trials, 40 V+-Congruent trials, 40 V+-Incongruent trials). At the beginning of the experiment, subjects performed two unisensory and one baseline session. During the formers, visual-(80 trials) and tactile-only (80 trials) stimulation was delivered, with subjects performing the same localization task as the MSI one described above. Importantly, visual and tactile stimuli were the same as the ones presented after each interaction. The latter would consist of 40 T+ and 40 V+ trials of the MSI task performed prior to any interaction, in order to let the participants getting familiar with the task and to measure cross-modal interference effects with no modulation related to the interactive session. The number of trials per condition was determined based on prior studies adopting similar paradigms targeting similar visuo-tactile neural processes (Sciortino et al., 2022).

### Electroencephalogram recording and processing

EEG signals were recorded and amplified using a gTec g.HIAMP Amplifier (g.Tec medical Engineering GmbH, Austria) and acquired from 126 active electrodes arranged according to the 10–10 system. Two electrodes were placed on the earlobes serving as reference, and all electrodes were physically referenced to the electrode placed on the right earlobe and then algebraically re-referenced off-line to the average of both earlobe electrodes. Impedance was kept below 30 KΩ for all electrodes for the whole duration of the experiment, amplifier hardware band-pass filter was 0.01-200 Hz and sampling rate was 1200 Hz. Offline, we applied a high pass filter to 0.1 and a low pass filter to 48 to remove slow drifts and line noise (50 Hz) from the signal. The two filters were applied separately as suggested by the latest version of EEGLAB (Delorme et al., 2004). Then, channels exhibiting excessive amounts of noise were detected by running the *clean_rawdata* function as implemented in EEGLAB (Delorme et al., 2004), by removing the channels which trace correlated with their direct neighbors for less than 80% of the time. After removing the above-mentioned channels, we epoched the continuous data into segments of 2400 milliseconds – from −1.200 to 1.200 around the onset of the stimulus. To identify and remove from the signal ocular artifacts (eye blinks and saccades) and excessive muscular noise, an Independent Component Analysis (ICA - Jung et al., 2000) was performed on the epoched data. Artefactual components were identified based on their time course and topography, and removed. On average, 9,9 (±2,75) components were removed per recording session (equivalent to 7,1% of the identified components per subject).

After the ICA, the bad electrodes previously removed (if present) were interpolated, with their signal being reconstructed based on the one from their surrounding neighbors with a “weighted” approach, as implemented in Fieldtrip (Oostenveld et al., 2011). Finally, EEG data were re-referenced to the average of all channels, and epochs containing remaining artefacts such as electrode jumps were removed following visual inspection.

### Data analysis

#### Behavioural Data

Reaction times (RTs) of the MSI task were taken as a measure of behavioural performance and interpersonal cross-modal integration effects. Only RTs from trials where subjects gave the correct response were taken into account for further analyses, with very few trials being inaccurate (average of inaccurate trials per subject = 2.4 ± 1.6%).

#### EEG Multivariate Analysis

We adopted a multivariate approach to explore whether the brain’s response to multisensory stimuli (i.e. multisensory processing) would change depending on the use of multisensory information and sensorimotor contingencies during a previous interpersonal interaction. Multivariate analyses assume that the processing of different stimuli has different neural patterns associated to be exploited. Here, we investigated whether the very same visuo-tactile stimuli would elicit different neural patterns depending on the observer’s previous interaction. Analyses were performed using the MVPA-Light toolbox (Treder, 2020) in Matlab R2022b and custom-made scripts. Prior to every classification, data were normalized with a 200 ms pre-stimulus baseline, downsampled to 500 Hz and z-scored to center and scale the training data, providing numerical stability (Treder, 2020). Moreover, if a class had fewer trials than another, we corrected the imbalance by under sampling the over-represented condition (i.e., randomly removing trials). Lastly, to increase the signal-to-noise ratio of the samples, training data (i.e. EEG trials) from the same class (i.e. M-PV, M-P, V) were split into groups of 5 and averaged.

#### Multiclass decoding of the interaction

First, we performed a multiclass decoding over time, investigating whether a Linear Discriminant Analysis (LDA) classifier was able to distinguish one condition (i.e. M-PV/M-P/V) from the other two based on the neural patterns in the time window from −200 to 1000 ms around stimulus onset. For linearly separable data, an LDA classifier divides the data space into *n* regions, depending on the number of classes, and finds the optimal separating boundary between them using a discriminant function to find whether the data fall on the decision boundary (i.e., 50% classification accuracy) or far from it (i.e., > 50% classification accuracy). A *k*-fold cross validation procedure in which samples were divided into 5 folds was repeated 20 times so that each sample was either used for training or testing at least four times, and the classifier’s raw output was visualized to assess decoding performance over time. Statistical significance (i.e., performance significantly above chance level) was assessed through non-parametric cluster-based permutation tests (Maris & Oostenveld, 2007), using ‘maxsum’ for cluster corrections and Wilcoxon test for significance assessment. As a control analysis, we performed the same multiclass decoding, respectively, on T+ -, V+ -, congruent- and incongruent-only EEG trials (see Supplementary Figure 1).

Secondly, following a purely data-driven approach, we looked at the multiclass LDA’s performance in a specific, early, time window to investigate whether the neural patterns differentiating the processing of multisensory stimuli after different interactions would be modulated already at low levels of the sensory processing cascade. To this aim, we performed a multiclass decoding, training and testing the classifier on the EEG data in the time window between stimulus onset (0 ms) and 150 ms. The accuracy values across testing folds of all repetitions were then averaged and presented on a confusion matrix to assess the probability of the classifier to in/correctly assign classes. As a control, we performed the same analysis on the EEG data in the time window between 150 and 300 ms post stimulus onset.

#### Binary decodings of the interaction

Third, we performed three binary classifications investigating how the performance of the classifier when distinguishing each condition from one of the other two evolved over time, as well as how stable (generalizable) the neural representations contributing to such decoding were. Specifically, we adopted the temporal generalization method (King & Dehaene, 2014), training the classifier on each single time point and testing it at all time points in the same time window previously selected for the multiclass classification over time (i.e., −200 1000 ms around stimulus onset). The generalization ability across time illustrates the similarity of EEG activity patterns relevant for encoding features and has been proposed to assess the stability of neural representations. Thus, if the previous interaction is uniformly encoded in the EEG activity patterns elicited by visuo-tactile stimuli (i.e., its influence is stable across time), then the performance of the classifier trained at time *t* will be shared to correctly decode it from EEG activity pattern at other time points (i.e., it would spread out of the diagonal in the time x time matrix of results). Vice versa, if the previous interaction transiently modulates distinct EEG activity patterns elicited by visuo-tactile stimuli (i.e., its influence is not stable across time, or it is limited to specific stages along the sensory cascade), then the performance of the classifier at time *t* will not generalize to other time points. This provided three temporal generalization matrices (time x time) with % accuracy values tested for statistical significance (i.e. performance significantly above 0.5 chance level) through non-parametric cluster-based permutation tests (Maris & Oostenveld, 2007). Moreover, to investigate what electrodes contributed most to these classifications over time, we performed three binary searchlight analyses using two neighboring matrices for time points and electrodes, respectively. Searchlight analysis is one approach to localize multivariate effects, as it strikes a balance between localization and statistical power (Kriegeskorte et al., 2006; Treder, 2020). Thus, in this analysis each electrode/time point and its direct neighbors acted as features for the classification, resulting in a channels × time points matrix of accuracy scores tested for statistical significance. By plotting the results from this matrix on a spatial topography we then visualized which electrodes carried the most weight in each temporal decoding.

#### Binary decodings of congruency and task relevance

Lastly, we investigated whether the neural patterns evoked by multisensory stimuli differed depending on 1) the congruency between the visual and the tactile information (i.e. decoding Congruency) or 2) whether subjects had to report the visual or the tactile information (i.e. decoding Task Relevance). Further, we also investigated the extent to which the LDA classifier generalized such differences across interactions (i.e. cross-decoding). In detail, we tested the temporal generalization of the neural patterns encoding Congruency or Task Relevance training and testing the classifier in the same condition (i.e., M-PV/M-P/V) as well as training it in one of the three conditions and testing it on the other two. This provided nine temporal generalization matrices (time x time) for each of the two classifications (i.e. Congruency and Task Relevance), with the three on the diagonal of the plot representing the within-condition decoding and the six off the diagonal representing the cross-condition decoding. As for the binary temporal generalization analyses classifying the previous interaction, these classification profiles highlighted how stable the neural patterns leading to the decoding were. Specifically, they showed if the mechanisms underlying (in)congruency or task-relevance processing were 1) non-generalizable (i.e., significant decoding accuracy only near the diagonal), thus indicating that such representations were transient, or 2) generalizable (i.e., significant decoding accuracy spreading outside the diagonal), thus indicating that such representations were sustained over time. Further, by cross-decoding these patterns, this approach allowed us not only to understand how Congruency and Task Relevance were decoded following each interaction, but also if the neural patterns differentiating such factors differed from one interaction to another, showing whether the brain forms (in)congruency or task-relevance representations that, at different stages, rely on neural generators whose activity is (or is not) modulated depending on a previous interpersonal interaction.

### Statistical Analysis

Linear mixed models were performed with R Studio software (2021), with single trials Reaction Times (RT) as the dependent variable. We included Condition (M-PV/M-P/V), Task Relevance (T+/V+) and Congruency (Congruent/Incongruent) as categorical predictors. Statistical significance of fixed effects was determined using type III ANOVA test with Satterthwaite’s method with the *lmer* function from *lme4* package (Bates et al., 2014) and the *anova* function. Thus, we modelled main effects for the Condition, Task Relevance and Congruency factors (both fixed and random effects) and the Participants as random intercept. We initially specified a maximal random effects structure (Barr et al., 2013), including random slopes for all fixed effects and their interactions by participant. However, due to non-convergence, we simplified the random-effects structure by including only the main effects as random slopes, which allowed the model to converge while still accounting for subject-level variability in the fixed effects. All tests of significance were based on an α level of 0.05. Normality checks were performed with the Shapiro-Wilk test. Post-hoc tests were performed using the Holm-Bonferroni method when appropriate via the *emmeans* (Lenth et al., 2019) packages. The *ggplots2* package (Wickham, 2016) was used to obtain boxplots of the data where mixed models analyses were performed.

## RESULTS

Overall, our results show that interpersonal motor interactions that rely on visual and/or motor-tactile/proprioceptive channels modulate both the behavioral (i.e., Reaction Times) and neural (i.e., evoked EEG patterns) markers of interpersonal cross-modal integration.

### Behavior

First, our 3 Condition (M-PV/M-P/V) × 2 Task Relevance (T+/V+) × 2 Congruency (Congruent/Incongruent) linear mixed model on Reaction Times (marginal R_2m_ = 0.22 and a conditional R_2c_ = 0.49) extend the classical individual cross-modal interference effect (main effect of Congruency [F(1, 10621) = 1615.69, p < .001]) of spatially incongruent stimuli to the case of interpersonal incongruency, with subjects being significantly slower when the visual and the tactile stimuli were spatially incongruent (i.e., a visual stimulus close to the confederate’s thumb was paired with a tactile stimulus on the subject’s index – Fig. 1B – or vice versa). Second, as already shown by previous studies on visuo-tactile perception (Göschl et al., 2014; Bresciani et al, 2006), subjects’ performance was overall better when reporting the location of visual (i.e., V+ condition) compared to tactile (i.e., T+ condition) information (main effect of Task Relevance [F(1, 23) = 50.78, p < .001]). Coherently with this visual dominance, although subjects’ performance was significantly worse both when reporting incongruent percepts focusing on the tactile (p < .001) or on the visual (p < .001) information, the interpersonal incongruency effect was stronger when reporting the tactile compared to the visual information (significant interaction between Congruency and Task Relevance [F(1, 10621) = 603.70, p < .001]), highlighting a stronger interferent effect of visual over tactile localization compared to the opposite (Fig. 2A).

**Figure 2.**
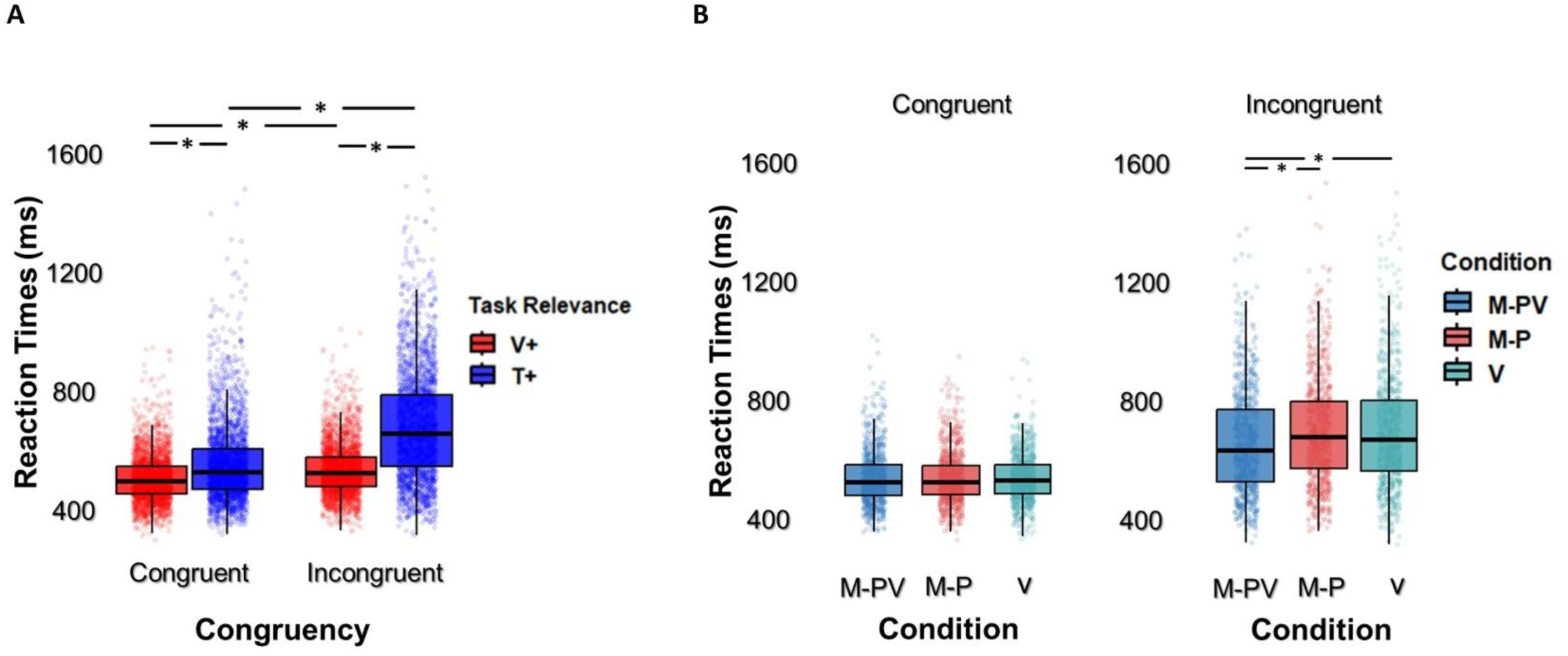
Behavioural results. A) Box plots showing the effect of spatial congruency (Congruent vs Incongruent) on the participants’ reaction times and its interaction with the Task Relevance factor. B) Box plots highlighting the facilitatory role of the M-PV interaction on the performance of the participants only when tactile location was task-relevant and a visual stimulus occurred in an incongruent configuration (right plot) compared to when the visual location was congruent with the tactile one (left plot). Each point represents one single trial; the central line in each boxplot represents the median; box limits represent upper and lower quartiles.

Crucially for the present study, a triple interaction between all the factors of the design emerged, highlighting that participants’ performance when focusing on touch, but not vision, and dealing with an interpersonal visuo-tactile incongruency was significantly modulated by the nature of the interpersonal interaction that preceded the cross-modal task (significant interaction between Condition, Task Relevance and Congruency factors [F(2, 10621) = 5.5, p = .004]). In details, subjects’ tactile reaction times in interpersonal incongruent trials with an interferent visual cue significantly diminished following the interaction involving both visual and tactile/proprioceptive modalities (i.e., the M-PV condition) compared to the two, single sensory modality, control conditions where either the visual (i.e., M-PV vs V; p = .04) or the motor-tactile/proprioceptive (i.e., M-PV vs M-P; p = .007) modality was used during the interaction (Fig. 2B). This pattern of results suggests an improvement in the processing of tactile stimuli independently from interferent visual stimuli (i.e., specific for incongruent trials) around the body of the partner after having interacted with them by using both visual, proprioceptive and motor modalities (i.e., higher interpersonal differentiation leading to a diminished cross-modal integration). This might be related to a modulation of the weights given to each sensory and motor modalities during the interaction that is later on reflected in optimal multisensory integration in incongruent trials (Ernst, 2002; Körding et al. 2007; Noppeney, 2021).

### Neural evoked responses

Our neurophysiological results complement the behavioural ones, showing a clear effect of different interpersonal interactions on the neural representations of interpersonal cross-modal integration. Indeed, the neural patterns evoked by (identical) visuo-tactile stimuli differed depending on the sensory channels involved in a previous interpersonal interaction. In detail, by adopting a multivariate approach, we show that a classifier was able to correctly identify the neural processing associated with trials following a specific interaction (multiclass decoding of the interaction) as early as 74 ms after the onset of the stimulus (Fig. 3B), independently from attentional factors (i.e., task relevance factor) or the congruency of visual and tactile stimuli (see supplementary results). Moreover, such an early modulation of neural responses appeared to be particularly dependent on the more ecological interaction (i.e., the M-PV one) where participants had to integrate information coming from their motor-tactile/proprioceptive and visual channels in order to fulfill the joint task with a confederate. Indeed, when comparing the decoding curves for the M-PV condition compared to the M-P and the V one, the performance of the classifier resulted significantly better in a time window starting as early as 110 ms after stimulus onset (significant clusters: M-PV vs M-P = [110 - 270 ms], *p*_clusterCorrected_ = .005; M-PV vs V = [140 - 416 ms], *p*_clusterCorrected_ = .007 - grey lines on top of Fig. 3B). Moreover, as shown in the confusion matrices in Fig. 3A, when relying only on very early (0-150 ms) evoked patterns of activity, the classifier’s capacity to distinguish the M-P from the V condition was worse than when it relied on later patterns (150-300 ms), while the M-PV trials were already clearly distinguishable in such an early time-window. This pattern strengthens the behavioral results, suggesting that the interpersonal interaction where vision and motor-tactile/proprioception were to be functionally integrated modulated differently the neural patterns related to the perception of following visuo-tactile stimuli.

**Figure 3.**
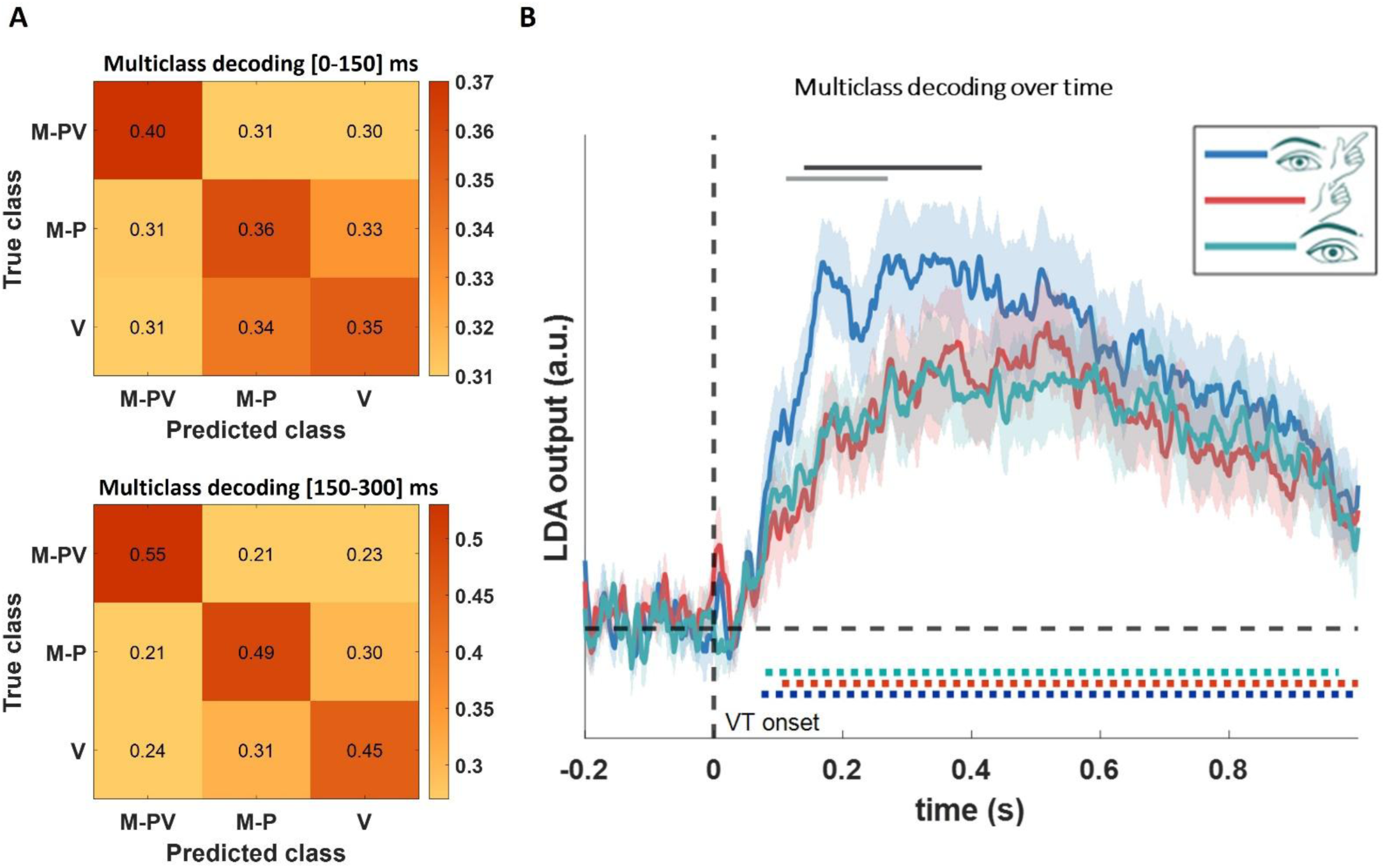
A) Confusion matrices showing the performance of the classifier in distinguishing the three behavioural interactions based on the neural patterns in the 0-150 ms time window (upper plot) and in the 150-300 ms time window (lower plot). B) Results from the multiclass decoding over time, showing that the classifier’s performance was significantly (*p*clusterCorrected < .05) above chance level for all three classes (dotted lines at the bottom), as well as highlighting the significantly (*p*clusterCorrected < .05) better performance of the classifier in distinguishing the neural patterns following the M-PV interaction compared to the ones following the M-P one (light-grey line on top), and to the ones following the V one (dark-grey line on top).

These results were complemented by the binary decoding of the previous interactions. These results allowed us to test i) how the performance of the classifier evolved over time when distinguishing each condition from one of the other two, ii) how stable (generalizable) the neural representations contributing to such decoding were, and iii) which electrodes carried the most weight to classification performance. The results of this approach are reported in Figures 4 and 5. The time × time matrices (Fig. 4), masked in order to show only significant results (*p*_clustercorrected_ < 0.05), provide two information complementing the first multiclass analysis. To begin with, they highlight three sustained generalization patterns, suggesting that the different modulation of the neural patterns evoked by visuo-tactile stimuli by the previous interactions was decodable in a sustained and generalized manner for each binary comparison from ∼200 ms up to 1000 ms following stimulus onset. Second, each decoding pattern reached statistical significance from ∼75 ms on, in line with the multiclass results. Interestingly, significant generalization blobs outside the diagonal occurring between ∼125 ms to 300 ms and before the sustained ones, starting from 200ms, were present only when classifying the M-PV condition compared to both the M-P and the V, but not when classifying the M-P compared to the V. This lack of generalization (i.e., non-significant off-diagonal areas between 125 and 300 ms) could reflect some early feedback of recurrent mechanism related to visuo-tactile processing in such specific time window which is specifically modulated by the M-PV interaction, leading to a defined and non-generalizable classification.

**Figure 4.**
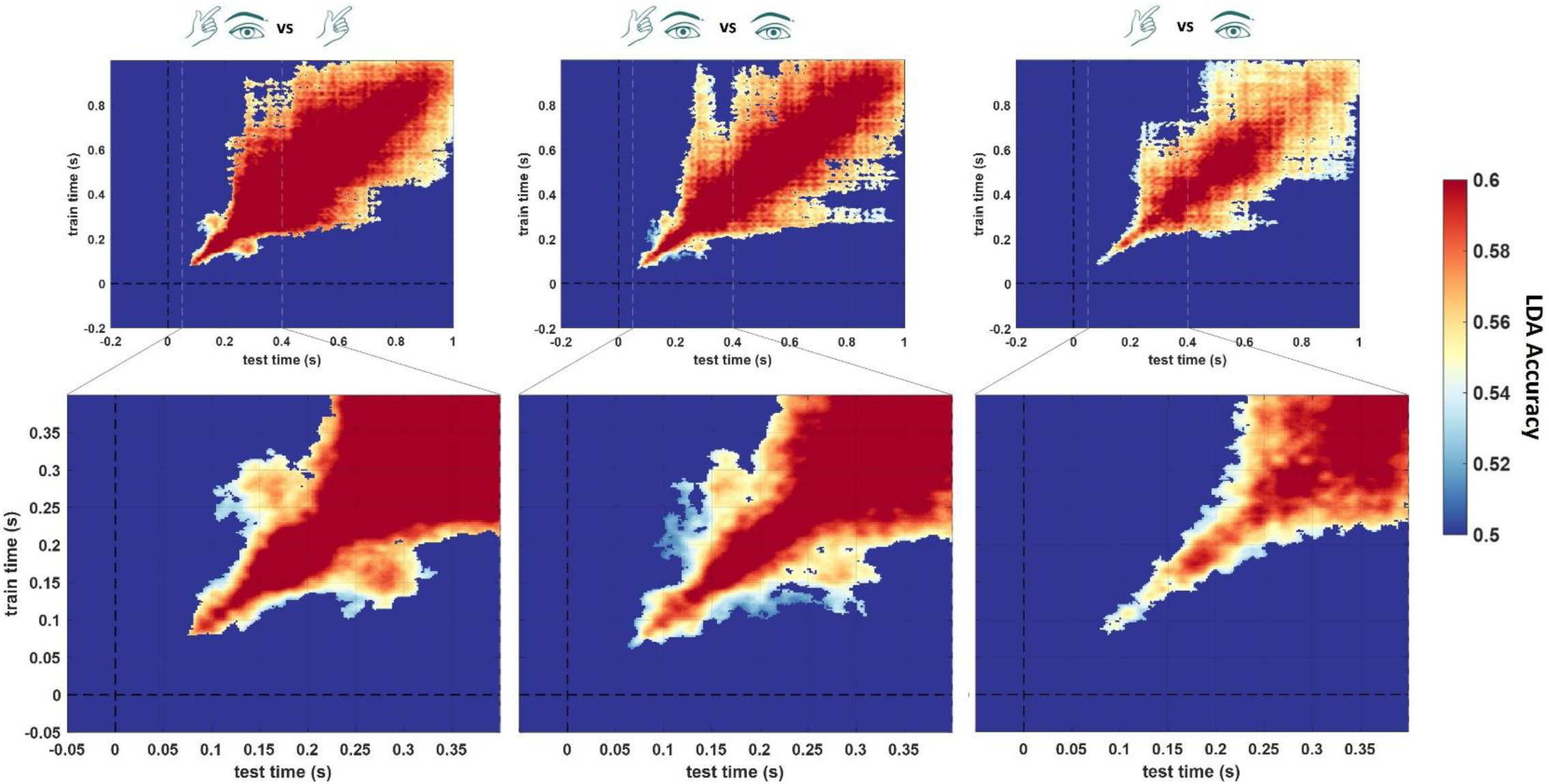
Results from the three binary classifications (i.e., M-PV vs M-P, M-PV vs V, M-P vs V). Temporal generalization matrices are masked in order to show only significant accuracy scores (pclusterCorrected < .05). White dashed lines highlight the early time-window (50 – 400 ms) following stimulus presentation presented in the bottom row plots.

**Figure 5.**
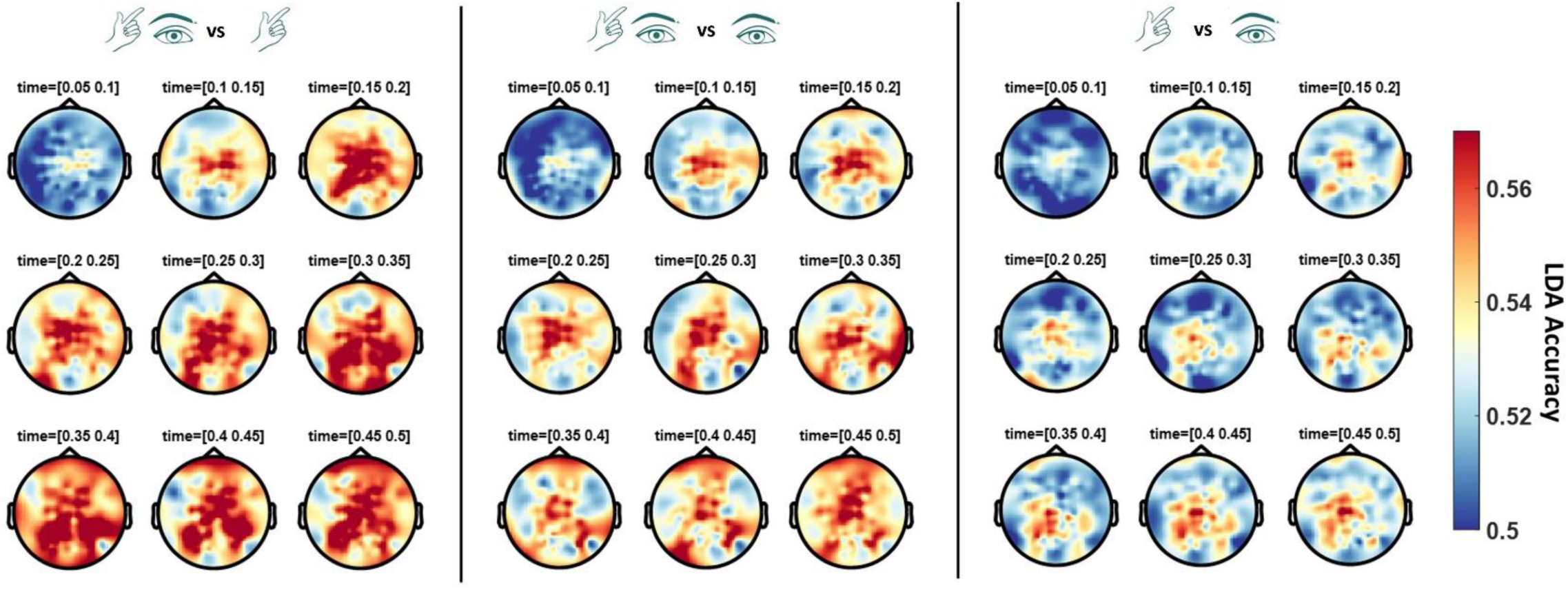
Searchlight analysis results plotted as topographies with accuracy values in a time-window between 50 and 500 ms following stimulus onset, separately for each binary decoding (MP-V vs M-P, MP-V vs V, M-P vs V).

The topographies shown in Figure 5 represent the weight of different electrodes to classification performance for each binary classification, obtained by running a searchlight analysis across time and electrodes with the same parameters and statistical analyses adopted for the classifications across time points only. Overall, the searchlight decoding is in line with the results across time, highlighting a sustained contribution of central, parietal and occipital electrodes to the classification of the electroencephalographic signal starting from 200 ms post-stimulus onset until the epoch’s end (1000 ms), which appeared to be reduced when classifying the M-P and the V conditions compared to when classifying the MP-V one. This highlights that the difference in visuo-tactile processing following the MP-V condition comes from a diffuse set of areas possibly where visual and tactile information are integrated.

The multiclass decoding as well as the binary decoding of the different interactions provided compelling evidence for a modulatory role of the interpersonal task on the processing of identical visuo-tactile interpersonal stimuli, irrespective of the congruency between the visual and the tactile information, or of the attentional focus (i.e., task-relevance) of each modality. To further investigate whether these perceptual characteristics changed depending on the previous interpersonal interaction, first we binarily decoded 1) congruent vs incongruent trials and 2) V+ vs T+ trials, separately for each condition. Second, we performed separate cross-decoding analyses to investigate the extent to which the Linear Discriminant Analysis (LDA) classifier generalized such binary classifications across different interpersonal interactions type. In detail, we tested the temporal generalization of the neural patterns encoding Congruency or Task Relevance training the classifier on one of the three interpersonal interaction conditions and then testing it on the other two. This analysis provides useful information on whether the neural information used by the classifier to decode Congruency or Task Relevance following one interaction would be as useful following a different one (i.e., cross-decodable), as well as how stable this cross-decoding would be (i.e., whether the decoding would be generalizable – relying on stable neural generators). The results from these analyses are plotted in nine temporal generalization matrices for each of the two factors (i.e., Congruency and Task Relevance), respectively in Figure 6 and 7. As shown on the three diagonal plots in Figure 6 the LDA classifier distinguished the neural patterns evoked by congruent compared to incongruent stimuli following all three interactions in a time window from 200 to 800 ms following stimulus onset. These classification profiles showed little to no generalization (i.e., significant decoding accuracy only near the diagonal), thus indicating that the representations of multisensory (in)congruencies were transient and not sustained over time. Most importantly, the LDA classifier was never able to generalize significantly better than chance from one condition to another before 400 ms (temporal generalization plots outside the diagonal, areas encircled by thick dark lines indicate significant generalization across interpersonal conditions), with the cross-decoding following the two conditions with the motor-tactile/proprioceptive channel involved (i.e., MP-V and M-P) sharing (in)congruency representations in a larger (thus, more sustained) time-window. In detail, the significant generalization areas decoding Congruency from MP-V to M-P and viceversa remain relatively stable from 400 to 800 ms, whereas both the one decoding Congruency from MP-V to V and viceversa and the one from M-P to V and viceversa share representations in two separate, short time windows, around 400 ms and around 700 ms. Overall, these cross-interaction generalizations suggest that at later stages (i.e., from 400 ms to 800 ms), the brain forms (in)congruency representations that rely on neural generators whose activity is not modulated depending on a previous interpersonal interaction. By contrast, the (in)congruency representations encoded in earlier (i.e., from 200 to 400 ms) EEG activity patterns did not enable cross-interaction generalization, suggesting that they are modulated by different interpersonal interactions.

**Figure 6.**
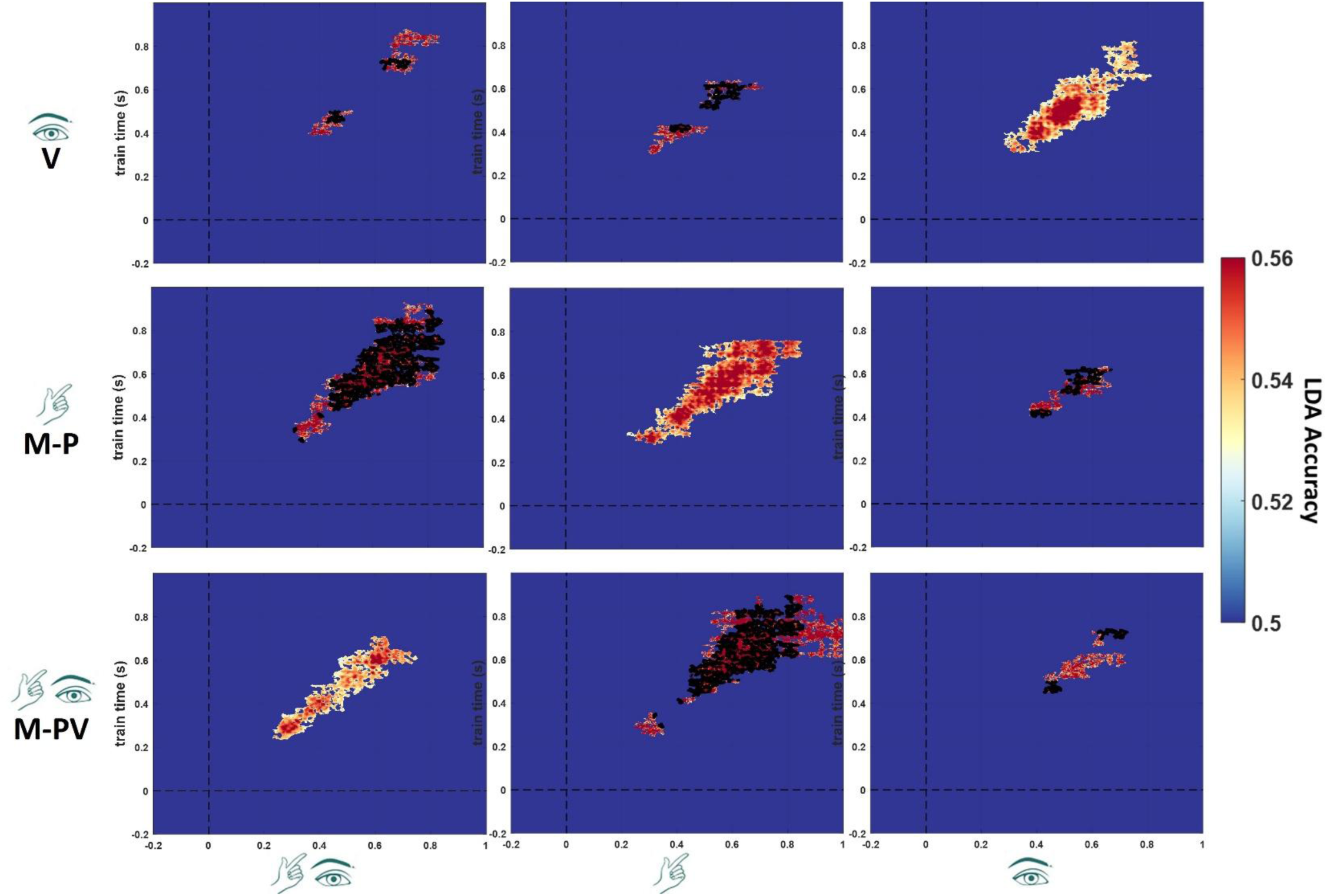
Results from the binary decodings and cross-decodings of Congruency (i.e., congruent vs incongruent trials). The three plots on the diagonal represent the decoding of Congruency training and testing the classifier on trials following the same interpersonal interaction. The six plots outside the diagonal represent the cross-decodings of Congruency, training the classifier on one of the three conditions, and testing it on the other two. All temporal generalization matrices are masked in order to show only significant accuracy scores (pclusterCorrected < .05). Areas encircled by thick dark lines encircle clusters with decoding accuracies that were significantly better than chance jointly for both 1) condition 1 to condition 2 and 2) condition 2 to condition 1 cross-temporal generalization.

**Figure 7.**
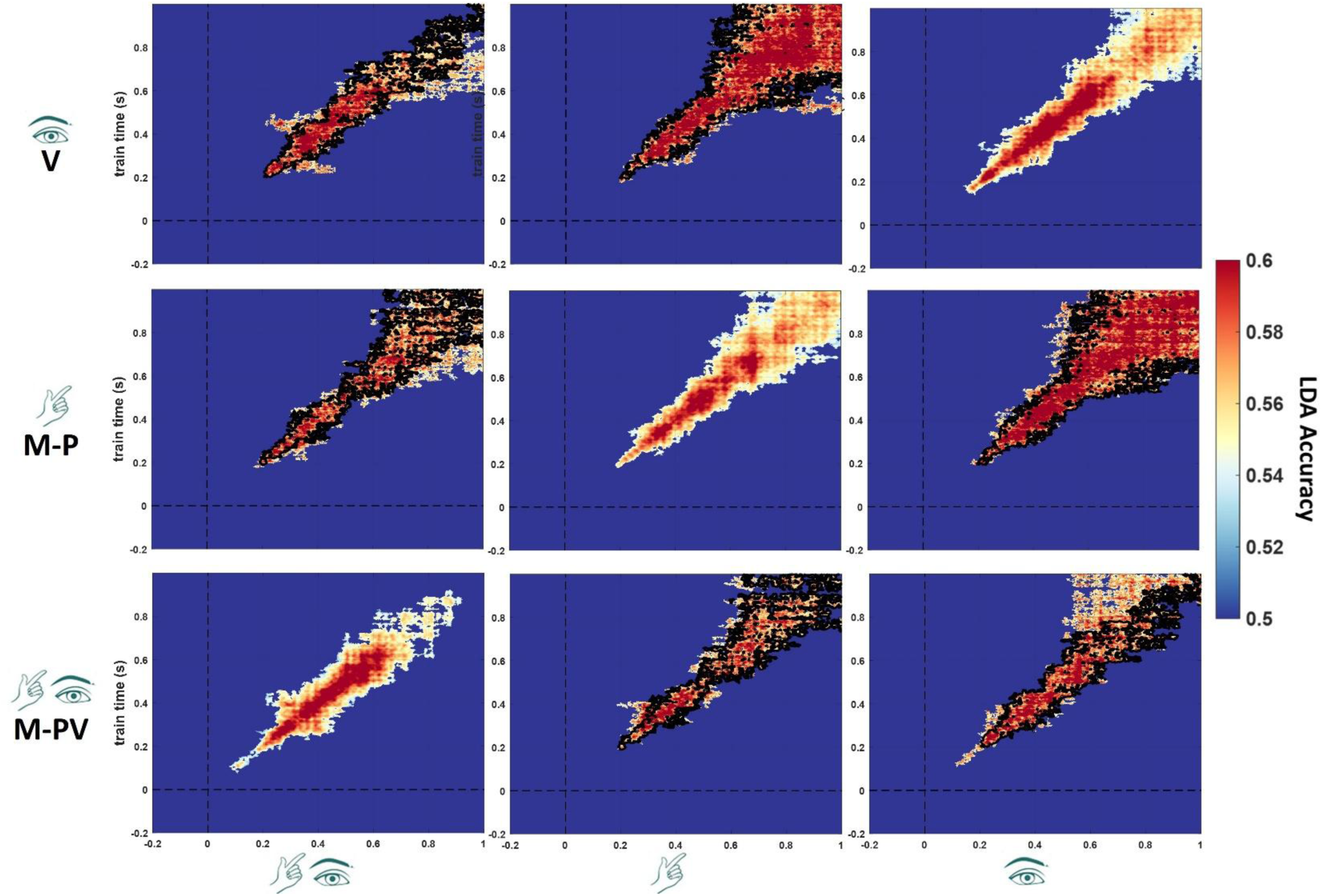
Results from the binary decodings and cross-decodings of Task Relevance (i.e., V+ va T+ trials). The three plots on the diagonal represent the decoding of Task Relevance training and testing the classifier on trials following the same interpersonal interaction. The six plots outside the diagonal represent the cross-decodings of Task Relevance, training the classifier on one of the three conditions, and testing it on the other two. All temporal generalization matrices are masked in order to show only significant accuracy scores (pclusterCorrected < .05). Areas encircled by thick dark lines encircle clusters with decoding accuracies that were significantly better than chance jointly for both 1) condition 1 to condition 2 and 2) condition 2 to condition 1 cross-temporal generalization.

The three diagonal plots in Figure 7 show that the LDA classifier distinguished the neural patterns evoked by visuo-tactile stimuli in the V+ compared to the T+ condition following all three interactions in a time window from 150 to 900-1000 ms following stimulus onset. These classification profiles showed little to no generalization (i.e., significant decoding accuracy only near the diagonal), thus indicating that the representations of top-down attentional processes were transient and not sustained over time. When cross-decoding across interactions, the LDA classifier was always able to generalize significantly better than chance from one condition to another from 200 ms onward (temporal generalization plots outside the diagonal, areas encircled by thick dark lines indicate significant generalization across interpersonal conditions). These cross-interaction generalizations suggest that the brain forms representations differing between V+ and T+ conditions that rely on neural generators whose activity is not modulated depending on a previous interpersonal interaction.

## DISCUSSION

In the present study, we recorded EEG in participants performing a multisensory spatial detection task to locate tactile stimuli that occurred together with somatotopically congruent or incongruent visual cues appearing near the hand of a partner with which they had performed different type of interactions. Specifically, we asked participants to actively engage in three interpersonal tasks where different sensory channels were recruited: in the Motor-ProprioVisual (M-PV) condition, subjects had to synchronize their movement with a partner while looking at their hand in order to reach a common goal (i.e., sawing a candle with a rope), thus relying on tactile/proprioceptive and visual information in order to hold tight the shared rope; in the Motor-Proprioceptive (M-P) condition, subjects performed the very same task with their eyes closed, thus focusing on the sensory information coming from their movement and from the rope in order to synchronize with the partner; and in the Visual (V) condition, subjects had to focus on the partner’s hand without moving, thus only focusing on the visual information obtained while monitoring the other’s movement. These interactions were designed to allow participants to use different sensorimotor channels to coordinate with a partner to test the hypothesis that these different interaction modalities would affect the way interpersonal intersensory information would be processed after the interaction.

To the best of our knowledge, the present results are the first ones in the literature showing that using different sensory modalities to achieve interpersonal coordination induces an offline modulation of both behavioral and neurophysiological markers of cross-modal interpersonal integration. Moreover, the present results extend previous literature on the influence of actions on sensory processes, where most of the studies focused on online effects occurring on a trial-by-trial basis (as the ones measuring adaptive responses to multisensory conflict - Noppeney, 2021). Furthermore, we extend previous results on the presence of cross-modal integration processes when sensory events happen close to one’s own and another person’s body, thus in an interpersonal (rather than individual) context. Indeed, Heed and colleagues (2010) showed that visuo-tactile interference effects are reduced in a cross-modal congruency task when visual and tactile stimuli appear, respectively, close to a confederate’s hand and on the participant’s hand, compared to when subjects performed the same task without the presence of the confederate in the peripersonal space (i.e., PPS – the space close around one’s body – Serino, 2019). These results complemented previous ones on the stronger integration of visual and tactile signals around the body (compared to far away – Pavani & Castiello, 2003; Spence et al., 2004) or around external objects that the brain presumably integrates into the body schema (i.e. rubber hands and tools – Pavani et al., 2000; Maravita et al, 2002). The authors interpreted these results as a proof of a top-down modulation of MSI in peripersonal space, which may contribute to guiding voluntary actions as well as to allow others’ actions in a space of self-defense. More recently, Fossataro and colleagues (2022) extended these results, observing multisensory enhancement with a novel setting developed in the azimuthal rather than in the radial direction (i.e., the other’s hand was positioned next to the participant’s one, rather than in front of it, as in the present study). The authors showed that the spatial proximity to someone else’s hand reduces the PPS boundaries, and that also multisensory EEG components were modulated by the proximity to one’s own or another person’s body. Altogether, these results have shown that visuo-tactile perceptual processes are modulated by the proximity of another person in the PPS, with an improvement in the processing of incongruencies (Heed et al., 2010) and a disruption of multisensory facilitation processes usually occurring around one’s own hand (Fossataro et al., 2022). Crucially, our results integrate these two approaches and show that visuo-tactile incongruencies are processed significantly better (i.e., they induce less interference) following an interaction with the confederate that required the visual and tactile/proprioceptive information to be integrated (i.e., the M-PV condition), compared to two interactions where either the visual (i.e. the M-P condition) or the motor-proprioceptive (i.e., the V condition) channels were absent. These results point towards a recalibration of the sensory weight of the visual or/and of the tactile-proprioceptive channel when previously integrated with vision, leading to a significant improvement in the processing of tactile information when a visual distracting stimulus is present. In other words, following an interpersonal motor interaction when vision and touch/proprioception were functionally integrated, participants showed less cross-modal interpersonal integration, i.e., a better self-other distinction. We speculate that the (multisensory) interaction with another agent contributing to the interpersonal motor exchange might lead either 1) to a defined representation of him/her as a different entity, or 2) to a better definition of one’s own body representation, or even 3) to a combination of the two, in turn leading to a weaker cross-modal interpersonal integration. Our results do not provide a clear evidence for one of these directions of effect, but pave the way for further studies comparing these hypotheses.

Most importantly, by leveraging MVPA, our study suggests a temporally precise modulation of the distributed cortical network involved in representing visuo-tactile stimuli (shared between one’s own hand and a partner’s hand) by different interpersonal motor interactions. In detail, the EEG responses evoked by identical visuo-tactile stimuli following different interactions were classified by a LDA classifier in a sustained manner lasting up to 1 second following their onset. Crucially, the responses following the M-PV interaction were clearly distinguishable from the ones following the M-P and the V condition in an early time-window (i.e., from 110 up to 400 ms), as shown by the multiclass decoding results. Binary decodings further extended this by showing that the neural generators whose activity was modulated by the previous interaction were activated in a sustained manner (i.e., the performance of the classifier generalizes more) in such an early time window when decoding the M-PV condition compared to the decoding of the M-P vs V conditions. Lastly, also the electrophysiological response to visuo-tactile spatial incongruencies differed based on the weight each sensory channel gained during the previous interaction. In detail, by cross-decoding the congruency factor from one condition to the other, we show that the EEG patterns around 200 ms to 400 ms encoding such stimulus characteristics were differently and uniquely modulated by each interpersonal interaction (i.e., no cross-interaction generalization was present), whereas at later stages (i.e., from 400 ms to 800 ms) the (in)congruency representations remained stable independently on the previous interaction. Interestingly, the evoked patterns related to top-down attentional processes (i.e., the decoding of the Task Relevance factor) were always cross-decoded from one condition to another. This suggests that higher-order computations during visuo-tactile spatial perception are not impacted by the re-weighting of sensory channels during a previous interaction. All in all, these results show that interacting with a confederate relying on the integration of different sensory channels impacts the early responses to visuo-tactile stimuli, including basic multisensory computations such as the processing of multisensory incongruencies. According to recent views on the development of body-part centered response fields (Bufacchi et al., 2025), these results could be interpreted in terms of a modulation of intersensory processing in the PPS (Ronga et al., 2021) depending both on social factors (Teneggi et al., 2013) and on the formation of different socSMC during the previous interpersonal interaction, leading to a re-weighting of each sensory channel.

To conclude, despite the evidence our results provide, we must acknowledge that the present paradigm focused on the performance of the subjects during the visuo-tactile spatial detection task, while we collected no measures regarding the interactive tasks. Further studies could extract, for example, the kinematic characteristics of the two partners moving, to be able to perform control analyses or even to analyze potential correlations between them and the following performance in the cross-modal task. Moreover, to obtain a full understanding of the way the interaction with the environment and/or other agents modulates the weights of different sensory modalities, further experiments across tasks (i.e., focusing on numerosity estimation; temporal discrepancies) and employing different analytical approaches (i.e., computational modelling) are needed.

In sum, the present results may pave the way for new approaches to study the multisensory foundations of interpersonal coordination, focusing on plastic and time-lasting changes in behavioral responses to sensory stimulation and neural representations of cross-modal effects depending on different contingencies (socSMC) between external sensory information (i.e., related to the environment and/or to other individuals) and internal motor and cognitive processing.

## Acknowledgments

We thank Matteo Gabucci and Cesare Gabucci for their contribution to the making of the custom-made wooden platform used during the interpersonal interaction task.

## Supporting information for

### Supplementary analyses and results

In order to check whether the classifier’s performance in the multiclass decoding over time (Fig. 3B) was different if we did not collapse across the Congruency and Task-Relevance factors, we performed the same analysis separately for T+, V+, congruent- and incongruent-only data. The results are plot in Figure S1, showing that, in all cases, the classifier’s performance in distinguishing the three classes was significantly above chance level. Moreover, the M-PV condition was always better classified than the other two classes (which, in turn, lead to similar classification patterns).

**Figure S1.**
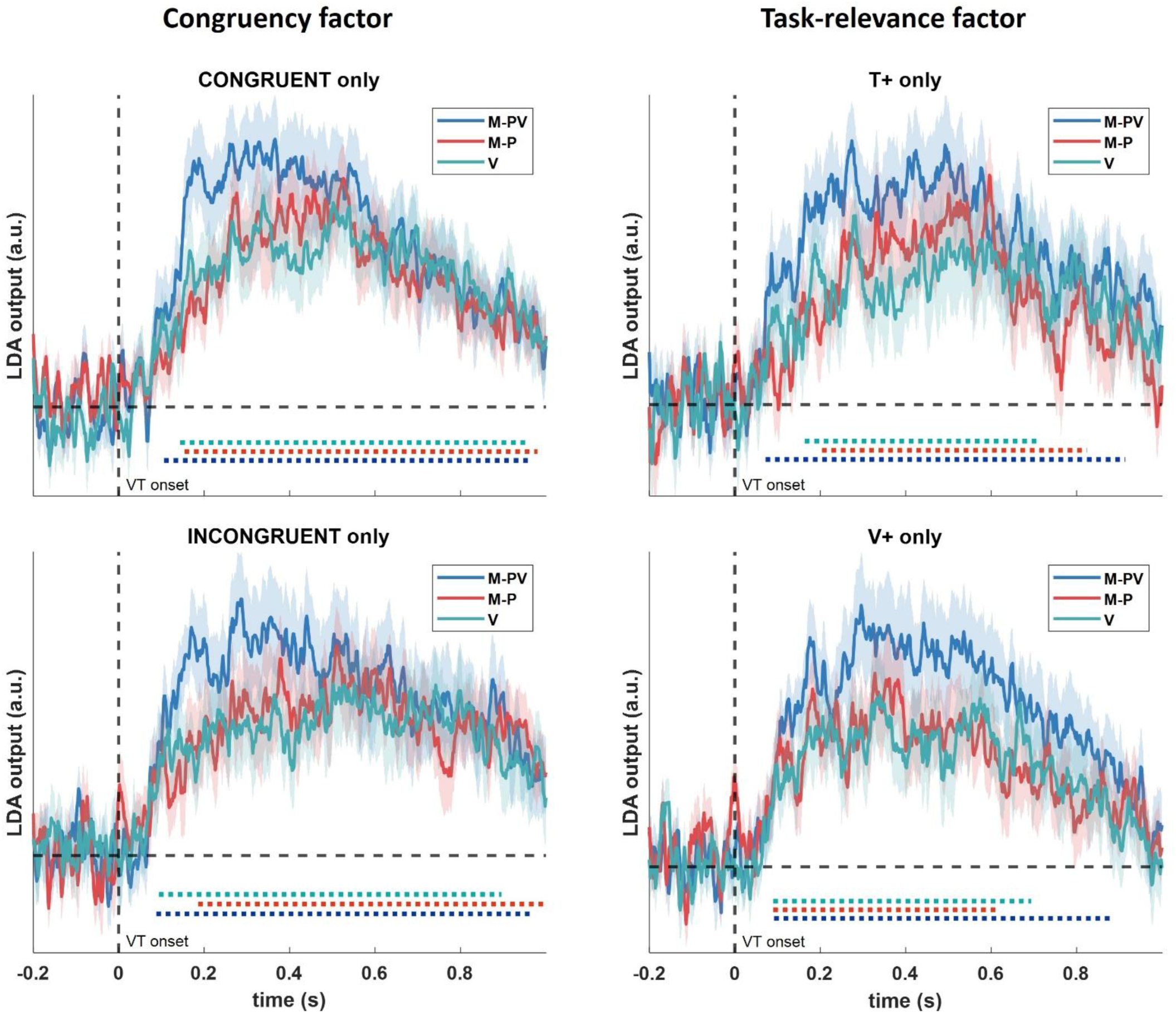
Results from the control multiclass classifications over time, with the time points when the classifier’s performance was significantly above chance level highlighted at the bottom, for each class. Left column: the classifier decoded the Condition separately for congruent and incongruent EEG trials. Right column: the classifier decoded the Condition separately for T+ and V+ EEG trials.

